# Somatic epigenetic mosaicism within *Pistacia terebinthus* (Anacardiaceae) trees leaves signature in reproductive cells

**DOI:** 10.1101/2025.11.14.688420

**Authors:** Carlos M. Herrera, Mónica Medrano, Conchita Alonso

## Abstract

Propagation of epimutations over the branching topology of individuals often produces epigenetic mosaicism in long-lived plants, but whether such epigenetic mosaics can bear some evolutionary relevance remains largely unknown. We test here whether epigenetic reprogramming during gametogenesis completely erases intraplant epigenetic variation or, alternatively, some “signature” of the somatic mosaicism persists into reproductive cells. Intraplant variation in global cytosine methylation of leaf and pollen DNA samples was studied in five adult males of the dioecious tree *Pistacia terebinthus* (Anacardiaceae). Paired samples of fresh expanding leaves and pollen grains were taken from five different twigs in each tree, and percent cytosine methylation estimated using HPLC. Trees were internally heterogeneous in global cytosine methylation of both leaf and pollen. Within-tree variance accounted for 49.7% and 30.5% of population-wide variance in methylation for leaves and pollen, respectively. Tree means and within-tree variances in pollen methylation were directly correlated, respectively, with tree means and within-tree variances in leaf methylation. As a consequence of the epigenetic variability accumulated in somatic tissues over the individual plants’ lifetimes being only weakly eroded by gametogenesis, intraplant epigenetic heterogeneity became a natural source of epigenetic gametic diversity in the wild-growing population of *P. terebinthus* trees studied.

## Introduction

Stable intraindividual mosaicism in DNA epigenetic features often arises in plants as a consequence of somatic epimutations (van der Graaf *et al*., 2015; Johannes, 2024) eventually propagating genealogically over the branching topology of individuals (Herrera *et al*., 2021; Yao *et al*., 2021, Yao *et al*., 2023). In addition, intraindividual epigenetic variation can be correlated with phenotypic variation across homologous structures of the same plant (Herrera & Bazaga, 2013; Alonso *et al*., 2018; Herrera *et al*., 2019, Herrera *et al*., 2022; Lloyd & Lister, 2022; Saiz-Blanco *et al*., 2025). These findings suggest that epigenetic mosaicism could contribute to the large intraindividual component of phenotypic variance ordinarily found in wild plant populations, which in turn has a number of ecological consequences for individuals (Herrera, 2009, 2017, 2024). For example, heterogeneity in global cytosine methylation within adult *Lavandula latifolia* Medik. (Lamiaceae) shrubs is related to intraplant variation in number and size of seeds produced per inflorescence (Alonso *et al*., 2018), and experimental augmentation of among-ramet heterogeneity in global cytosine methylation amplifies intraplant variance of vegetative and reproductive traits in the perennial herb *Helleborus foetidus* L. (Ranunculaceae) (Herrera *et al*., 2019). Limited evidence also suggests that epigenetic mosaicism within plants can have transgenerational effects by enhancing variance in growth and survival of progenies from epigenetically dissimilar modules of the same plant (Herrera *et al*., 2022). For these effects to have some evolutionary significance, however, somatic epigenetic heterogeneity acquired over a parent plant’s lifetime via patchy environmental induction or random epimutations (Herrera *et al*., 2021; Yao *et al*., 2021, Yao *et al*., 2023; Shah, 2022; Gardner *et al*., 2023; Chan *et al*., 2024; Vo *et al*., 2024) should translate into epigenetic and/or phenotypic differences in its adult progeny.

Obtaining this crucial evidence with traditional heritability estimation methods is impractical, given the long lifespans of plants where epigenetic mosaicism should be most frequent and, possibly, evolutionarily relevant too. One feasible alternative is to examine whether the somatic epigenetic mosaicism in vegetative, diploid tissues of the parent plant is at least partly preserved in post-gametogenesis, haploid reproductive tissues (ovules or pollen). The rationale for this approach stems from the fact that thorough epigenetic reprogramming, including changes in frequency and genomic distribution of cytosine methylation, often takes place during gametogenesis, the process that connects the sporophytic and gametophytic generations (Gutierrez-Marcos & Dickinson, 2012; Kawashima & Berger, 2014; Jo & Nodine, 2024). Intraplant epigenetic variation accumulated over an individual’s lifetime could therefore be wiped out during gametogenesis, leading to all plant parts eventually producing epigenetically homogeneous pollen or ovules. Consequently, a necessary (albeit not sufficient) requisite for intraplant somatic epigenetic mosaicism possessing some transgenerational significance is that epigenetic reprogramming during gametogenesis were insufficient to erase intraplant epigenetic heterogeneity, so that some discernible “signature” of such somatic heterogeneity would persist past gametogenesis. In other words, that pollen or ovules produced by epigenetically distinct parts of the same adult plant should be also epigenetically heterogeneous. This paper tests this possibility for male trees of *Pistacia terebinthus* L. (Anacardiaceae) by comparing intraplant variation in global DNA cytosine methylation in vegetative (leaves) and reproductive tissue (pollen) across branches of the same tree.

## Materials and methods

### Study plant and field methods

*Pistacia terebinthus* is a small deciduous, dioecious, wind-pollinated tree widely distributed in the Mediterranean Basin. In male trees, clusters of fresh leaves develop in spring at the end of twigs at the same time that nearby inflorescences mature and shed pollen. Simultaneous timing and closeness of leaves and inflorescences on the same twig make possible paired sampling of somatic and reproductive tissues of identical age in the same growth axis. *Pistacia terebinthus* trees are long-lived and branch out profusely from one or a few trunks rising from a single basal rootstock. Reiteration of the paired sampling scheme across different twigs in the same crown permits a concurrent assessment of within-plant variability of DNA cytosine methylation in fresh leaves (i.e., diploid somatic cells) and pollen (i.e., post-gametogenesis haploid reproductive cells) while controlling for possible seasonal or age-related differences in DNA methylation. Another advantage of the *P. terebinthus* study system is that male inflorescences produce copious pollen, which facilitates obtaining sufficient pollen DNA for chemical analysis of post-gametogenesis cytosine methylation levels (see below).

Five single-trunk, male *P. terebinthus* trees (trunk perimeter 40 cm above ground = 33-84 cm) were chosen for study in a dolomitic outcrop at 1100 m a.s.l. in the Sierra de Cazorla, Jaén province, Spain. Within each tree, five paired samples each consisting of expanding leaves and pollen-shedding inflorescences in the same twig were collected from twigs as widely spaced as possible within the crown. Leaf samples were placed inside paper bags, dried at ambient temperature within containers filled with silica gel, and kept at ambient temperature until processed. To promote anther dehiscence and pollen shedding, inflorescence samples were left to dry on aluminium foil sheets within containers with silica gel. Once anthers had released most pollen the aluminium foil piece was closed by folding it over itself, stored in a paper bag and kept dry at ambient temperature until processed.

### Laboratory methods

Dry leaf samples were homogenized to a fine powder using a Retsch MM 200 mill. Pure pollen samples were obtained by separating pollen grains from inflorescence remains (anther walls, inflorescence axes, bracts) following with minor modifications the procedure of Johnson-Brousseau & McCormick (2004), based on using different-pore sized Nitex® meshes (80, 38 and 6 µm pore size meshes in our case; Sefar America, Inc., Depew, NY, USA), held together in sequence and attached to a handheld vacuum cleaner. The 80 µm mesh filtered out most of the larger flower parts and debris of unwanted maternal tissue, while the 38 µm mesh trapped smaller debris and small amounts of pollen. Most of the pollen passed through these two first mesh filters and was trapped on the third, 6 µm mesh. Samples of dry material stored into the aluminium foil bags were released inside hand-made 80 µm mesh bag and immediately placed inside other two nested mesh bags (38 µm and 6 µm for inner and outer bags, respectively). To harvest the purified pollen sample, the set of three nested bags with decreasing pore size holding the plant material was placed inside a plastic plumbing tube attached to a handheld vacuum cleaner and exposed to the suction force of the device. A paintbrush was used to release the pollen gathered into the 6 µm filter, which was stored dry into a 1.5 mL microcentrifuge tube. This procedure was repeated on aliquots of the same field sample until sufficient quantity of pollen was obtained. On average, 125 mg of purified pollen was obtained from each field sample (range = 40-330 mg).

Total genomic DNA was extracted from leaf (*N* = 25) and pollen (*N* = 25) samples using the Omega Biotek E.Z.N.A. Plant DNA DS Mini Kit, and digested with DNA Degradase Plus (Zymo Research), a nuclease mix degrading DNA to its individual nucleoside components. Digested samples were stored at −20°C until analysed. To allow for properly testing the statistical significance of within-tree heterogeneity (i.e., among twigs) in DNA methylation level of leaves and pollen, two (leaves) or six (pollen) aliquots were prepared from each DNA hydrolyzate, and the resulting *N* = 200 samples (5 plants x 5 twigs x 8 aliquots) were processed in randomized order. Genome-wide percent methylation of DNA cytosines was determined for each sample with the chromatographic technique described in detail by Alonso *et al*. (2016), using reversed phase HPLC with spectrofluorimetric detection. Global cytosine methylation was estimated for each sample as 100 x 5mdC/(5mdC + dC), where 5mdC and dC are the integrated areas under the peaks for 5-methyl-2’-deoxycytidine and 2’-deoxycytidine, respectively. The position of each nucleoside was determined using commercially available standards. Samples yielding poor-quality or insufficient DNA, or producing weak chromatographic signal and unreliable peak area integration, were discarded. Results presented below are based on 45 (leaves) and 143 (pollen) independent HPLC runs (see Herrera 2025 for raw data).

### Statistical analyses

Statistical analyses reported in this paper were carried out using the R environment (version 4.5.0; R Core Team, 2025). Statistical significance of heterogeneity among twigs of the same tree in percent cytosine methylation of leaf and pollen DNA was tested by fitting separate linear fixed-effects models to leaf and pollen data using the lm function in the stats package, with tree and twig nested within tree as predictors. The statistical significance and magnitude of DNA methylation differential between leaves and pollen from the same twig were assessed by fitting a linear mixed-effects model to percent methylation data, with plant tissue (leaf *vs*. pollen) as fixed effect, and tree and twig nested within tree as random effects, using the lmer function in the package lme4 (Bates *et al*., 2015). Hierarchical partitions of total sample-wide variance in percent cytosine methylation of leaf and pollen DNA into components attributable to tree, twig nested within tree and residual measurement error, were conducted by fitting intercept-only linear mixed-effects models to data with the lmer function. In all fitted models the statistical significance of predictors was determined with log-likelihood tests.

## Results

### DNA methylation in leaves

Percent DNA cytosine methylation of leaf tissue varied significantly both among trees and among twigs within the same tree, thus revealing epigenetic heterogeneity of vegetative plant parts in the sampled population at both the individual and intraindividual levels (Table 1, Figure 1). The trees sampled differed widely in among-twig variability of leaf DNA methylation, which was highest in tree P1 (methylation range = 14.6–17.0 %), lowest in tree P2 (methylation range = 15.8–16.1%), and intermediate in the rest (Figure 1).

**Table 1.**
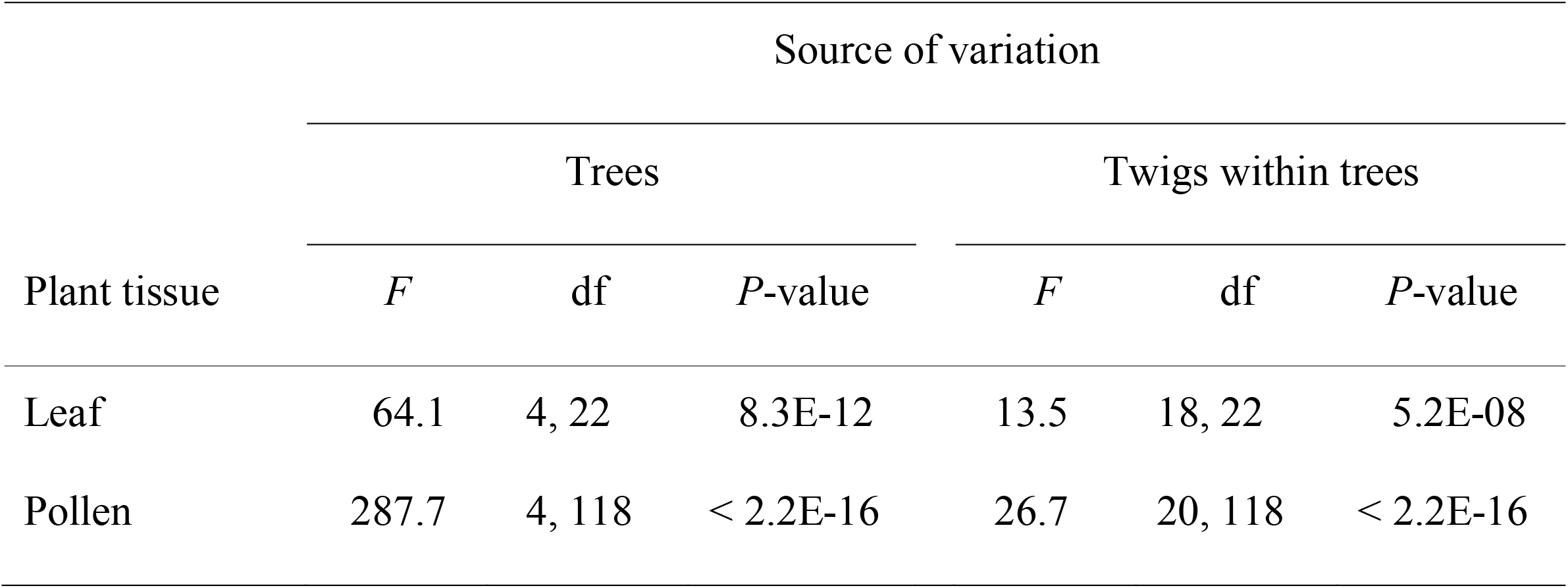
Summary of results of linear fixed-effects models testing for significance of heterogeneity among trees, and among twigs within trees, in percent global DNA cytosine methylation of leaf and pollen sampled from the same twigs of *Pistacia terebinthus* trees following a paired design (*N* = 5 trees, 5 twigs per tree).

| Plant tissue | Source of variation |  |  |  |  |  |
| --- | --- | --- | --- | --- | --- | --- |
|  | Trees |  |  | Twigs within trees |  |  |
|  | *F* | df | *P*-value | *F* | df | *P*-value |
| Leaf | 64.1 | 4, 22 | 8.3E-12 | 13.5 | 18, 22 | 5.2E-08 |
| Pollen | 287.7 | 4, 118 | < 2.2E-16 | 26.7 | 20, 118 | < 2.2E-16 |

**Figure 1.**
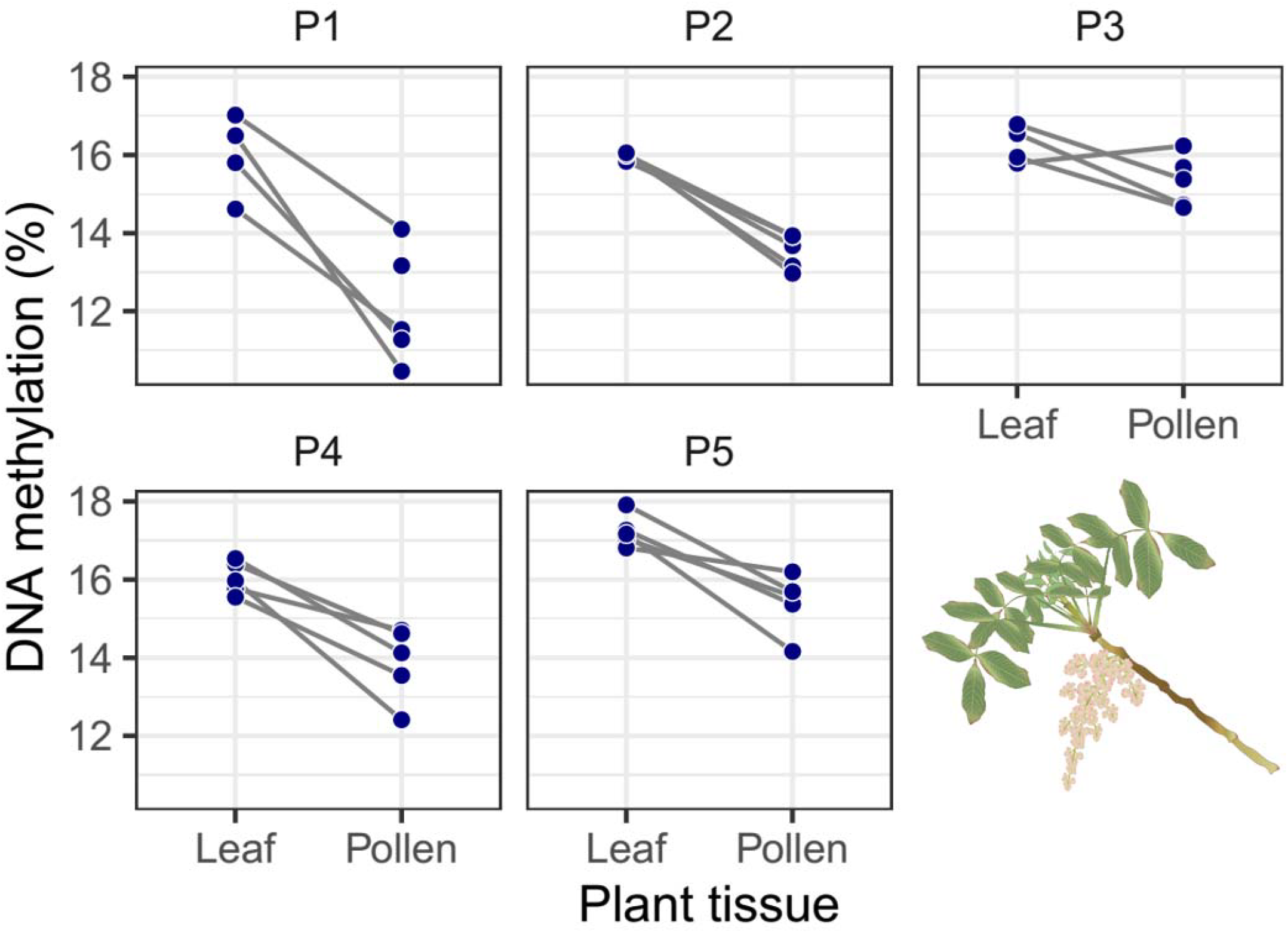
Percent global DNA cytosine methylation in leaf and pollen of the five trees of *Pistacia terebinthus* sampled (P1-P5). Dots represent the average methylation of all replicates for each tree x twig x tissue combination. Each line connects the leaf and pollen values for one twig in the same tree, hence vertical spread of lines for a given tree denotes within-tree heterogeneity in DNA methylation level in leaves and pollen. Inset shows a schematic depiction of the extreme of a *Pistacia terebinthus* twig at leaf flushing, showing the disposition of expanding leaves and pollen-shedding inflorescences.

Variance partitioning of total sample-wide variance in percent DNA methylation of leaf tissue revealed that differences between twigs of the same tree were quantitatively more important (49.7% of total variance) than differences among trees (42.4% of variance), thus denoting a remarkable importance of somatic epigenetic heterogeneity within trees.

### DNA methylation in pollen

There also existed statistically significant DNA methylation heterogeneity in the pollen, both among trees and among twigs of the same tree (Table 1), thus indicating that the methylation level of the DNA in the pollen grains produced by a given individual tree depended on which was the twig which produced it. As with leaf tissue, trees also differed in the magnitude of among-twig differences in pollen DNA methylation, the range being again broadest in tree P1 (methylation range = 10.5–14.1 %), narrowest in tree P2 (methylation range = 13.0– 13.9%), and intermediate in the rest (Figure 1).

The contributions of differences among trees, and among twigs of the same tree, to sample-wide total variance in DNA methylation of pollen were 30.5% and 62.9%, respectively. About one third of total population-wide variance in pollen DNA methylation was therefore due to differences among twigs of the same tree. Compared to the results of variance partitioning for leaf DNA methylation shown in the preceding section, these figures reveal a slightly weaker, but still substantial contribution of among-twig variation to population variance in pollen DNA methylation.

### Comparative leaf and pollen DNA methylation

Percent DNA methylation was consistently lower in pollen than in leaves of the same twig (Figure 1). The mean (± SE) leaf-to-pollen differential in percent methylation within the same twig averaged −2.07 ± 0.15%, denoting a measurable but small net genomic demethylation in the transition from somatic to reproductive cells (Chi-squared = 180.8, df = 1, *P* < 2.2E-16; mixed-effects model with plant tissue as fixed effect, and tree and twig nested within tree, as random effects).

Tree means and within-tree variances in pollen DNA methylation were positively correlated, respectively, with tree means and within-tree variances in leaf methylation (*r* = 0.695 and 0.904, *N* = 5, *P* = 0.016 and 0.036, for means and variances respectively; significance levels assessed using permutation). There was a positive, statistically nonsignificant relationship across twigs of the same tree between percent methylation of pollen and leaves (model parameter estimate ± SE = 0.54 ± 0.38, *P* = 0.14; mixed-effects model with pollen methylation as response, leaf methylation as predictor, and tree as random effect).

## Discussion

The recent discovery of a fast-ticking evolutionary epigenetic clock in plants has opened unexplored research avenues at the interface between ecology and evolutionary biology (Yao *et al*., 2023). Unfortunately, progress in ecological epigenetics (*sensu* Bossdorf *et al*., 2008) has been hindered by the methodological problems associated with epigenetic research on non-model wild plants lacking genomic resources. One of the alternative approaches developed for circumventing this dificulty is the study of variation in genome-wide levels of DNA cytosine methylation, which allows the characterization of individuals, populations and species from an ecological epigenetics perspective (Alonso *et al*., 2016, Alonso *et al*., 2018, Alonso *et al*., 2019; Agrelius *et al*., 2018; Singh *et al*., 2024). Changes in cytosine methylation in plants are involved in many biological processes, and variation in genome-wide methylation levels reflects the dynamic balance between methylation gain and loss in response to internal (ontogenetic) and external (ecological) signals, and can be heritable (Herrera *et al*., 2018; Liu *et al*., 2023; Cao & Chen, 2024; Kumar *et al*., 2024; Alonso *et al*., 2025). These features justify using genome-wide methylation level for describing epigenetic variation at the within- and among-individual level as done here (see also Gao *et al*., 2010; Agrelius *et al*., 2018; Singh *et al*., 2024).

Sampled trees were internally heterogeneous with regard to global DNA cytosine methylation of leaf tissue. Internal divergence within trees was extensive, as within-tree variance accounted for half (49.7%) of total sample-wide variance. This within-plant variability in DNA cytosine methylation of leaves does not seem exceptional, as a similar value was reported for the shrub *Lavandula latifolia*, where leaf methylation differences among modules of the same individual accounted for 50.3% of population-level variance (Alonso *et al*., 2018). Significant epigenetic reprogramming took place during gametogenesis in *P. terebinthus*, involving a net average demethylation of −2% relative to the methylation level of leaf tissue (≈ % *de novo* methylation minus % demethylation).

Despite this reprogramming, however, individual trees still remained heterogeneous in the methylation level of pollen produced by different twigs, and such variation accounted for 30.5% of sample-wide variance instead of the near-zero component one would expect from any efficacious, thorough epigenetic reprogramming over gametogenesis. This result supports the view that epigenetic mosaicism of homologous structures within adult plants can involve both vegetative and reproductive tissues, as implied by the epigenetic mosaicism hypothesis of intraplant variation (Herrera *et al*., 2021). We are not aware of other similar investigations considering global cytosine methylation of diploid vegetative and haploid reproductive tissues in non-model plants in the wild.

In adult long-lived plants such as *P. terebinthus*, intraindividual heterogeneity in DNA methylation of vegetative tissues is most likely the combined consequence of an epigenetic molecular clock that continuously generates random variability (Herrera *et al*., 2021; Yao *et al*., 2021, Yao *et al*., 2023; Gardner *et al*., 2023; Vo *et al*., 2024) and localized epigenetic changes induced by ecological factors impinging differentially on different sectors of a tree (e.g., herbivory, light regime, temperature, insolation; Roslin *et al*., 2006; Herrera & Bazaga, 2013; Zonner & Renner, 2015; Eisenring *et al*., 2021; Emmerson *et al*., 2025; Proβ *et al*., 2025). The intraplant epigenetic heterogeneity thus generated can underlie intraplant phenotypic heterogeneity with short-term ecological consequences for the individual, but these aspects will not be discussed here. Internal epigenetic heterogeneity of adult plants could also translate into longer-term effects such as, for instance, enhancing progeny heterogeneity in ecologically relevant traits, a possibility that remains little explored to date. Although epigenetic reprogramming during fertilization (Gutierrez-Marcos & Dickinson, 2012; Jo & Nodine, 2024) could further erode intraindividual variability in pollen methylation level found here, it seems unlikely that this additional reprogramming would be strong enough to entirely wipe out the epigenetic variation among pollen of different twigs in the same tree which “survived” the reprogramming during gametogenesis. Our results for *P. terebinthus* therefore provide plausible evidence that intraindividual epigenetic heterogeneity in vegetative parts may cascade into subsequent generations via epigenetically heterogeneous gametes. This implies that some epigenetic variants arising in somatic tissues over an individual plants’ lifespans will eventually make part of the set of epigenetic variants represented in the plant’s gametic pool and potentially contributing to the phenotypic diversity of the next generation (Shahzad *et al*., 2025).

The trees sampled differed widely in the extent of within-plant variation in cytosine methylation of leaf tissue. Causal factors accounting for differences between trees can only be speculated at present, but we suggest that they should partly reflect differences in age, since under a random epigenetic clock trees should become more epigenetically variable as they get older (Herrera *et al*., 2021; Lesaffre, 2021). Irrespective of the mechanisms involved, however, a remarkable result of our study is that, across trees, intraindividual variability in pollen DNA methylation closely mirrored intraindividual variability in leaf

DNA methylation. This result has implications that point to novel pathways for epigenetic research on long-lived plants. First, epigenetic reprogramming during gametogenesis not only failed to completely erase intraplant variation in leaf methylation level, but it was also unable to either blur or shrink the broad among-tree differences in within-tree variability. Second, transgenerational fecundity effects of intraplant epigenetic heterogeneity in vegetative tissue (Herrera *et al*., 2019, Herrera *et al*., 2022) could be mediated by the latter’s influence on epigenetic heterogeneity of gametes. Third, intraplant epigenetic heterogeneity of pollen could ultimately underly the intraplant variation in pollen grain size and viability found in some plants (Strelin *et al*., 2025). And fourth, the inability of epigenetic reprogramming during gametogenesis to distort or wipe out individual differences in intraplant epigenetic variability levels should generate an opportunity for selection on genes which limit or prevent the transgenerational transmission of environmentally-induced epigenetic states (e.g., Iwasaki & Paszkowski, 2014).

Dismissal of the possible evolutionary significance of intraindividual phenotypic variation among homologous reiterated structures produced by the same plant can be traced back at least to Wilhelm Johanssen’s (1911) distinction between genotype and phenotype, and his emphasis on the transgenerational irrelevance of phenotypic variation within isogenic lines (Mayr, 1982). Ironically, a recent re-analysis of Johanssen’s experimental results showed that phenotypic variation within pure genetic lines did actually have transgenerational phenotypic consequences (Herrera, 2024). In addition, as long as intraindividual phenotypic variation in long-lived plants has some stable epigenetic basis, the recent advances in the understanding of the molecular mechanisms accounting for the appearance and subsequent transgenerational preservation of epigenetic variants in plant populations (Stajic & Jansen, 2021; Yao *et al*., 2021, Yao *et al*., 2023; Fitz-James & Cavalli, 2022; Shahzad *et al*., 2025) should also prompt for a critical reassessment of the traditional dismissal of intraplant variation as a possible evolutionary force in plants. Our finding here that intraplant epigenetic heterogeneity in somatic tissues can translate into concomitant heterogeneity of the male gamete populations further stresses the necessity for such reconsideration.

## Ethics

This work did not require ethical approval from a human subject or animal welfare committee.

## Data availability statement

Raw data used in the study are available at figshare (Herrera 2025).

## Declaration of AI use

We have not used AI-assisted technologies in creating this article.

## Author contributions

CMH designed the research, conducted field sampling, performed statistical analyses and led the writing; MM developed the pollen purification protocol and prepared pollen samples; CA supervised HPLC analyses, curated results and contributed to interpretations. All authors refined the manuscript and approved the final version.

## Acknowledgements

We thank Pilar Bazaga and Mónica Gutiérrez-Rivillo for laboratory assistance, and Consejería de Medio Ambiente, Junta de Andalucía for permission to work in the Sierra de Cazorla and providing invaluable facilities there.

